# Vocal tract allometry in a mammalian vocal learner

**DOI:** 10.1101/2021.10.29.466455

**Authors:** Koen de Reus, Daryll Carlson, Alice Lowry, Stephanie Gross, Maxime Garcia, Ana Rubio-Garcia, Anna Salazar-Casals, Andrea Ravignani

**Author notes:** These authors share joint first authorship (Koen de Reus, Daryll Carlson).

## Abstract

Acoustic allometry occurs when features of animal vocalisations can be predicted from body size measurements. Despite this being considered the norm, allometry sometimes breaks, resulting in species sounding smaller or larger than expected for their size. A recent hypothesis suggests that allometry-breaking mammals cluster into two groups: those with anatomical adaptations to their vocal tracts and those capable of learning new sounds (vocal learners). Here we test which mechanism is used to escape from acoustic allometry by probing vocal tract allometry in a proven mammalian vocal learner, the harbour seal (*Phoca vitulina*). We test whether vocal tract structures and body size scale allometrically in 68 young individuals. We find that both body length and body mass accurately predict vocal tract length and one tracheal dimension. Independently, body length predicts vocal fold length while body mass predicts a second tracheal dimension. All vocal tract measures are larger in weaners than in pups and some structures are sexually dimorphic within age classes. We conclude that harbour seals do comply with anatomical allometric constraints. However, allometry between body size and vocal fold length seems to emerge after puppyhood, suggesting that ontogeny may modulate the anatomy-learning distinction previously hypothesised as clear-cut. We suggest that seals, like other species producing signals that deviate from those expected from their vocal tract dimensions, may break allometry without morphological adaptations. In seals, and potentially other vocal learning mammals, advanced neural control over vocal organs may be the main mechanism for breaking acoustic allometry.

## Introduction

In many species, acoustic signals help mediate social interactions such as competition for mates and territory, and parent-offspring recognition (Bradbury and Vehrencamp, 1998; Martin et al., 2017). Signals can encode information about the caller’s biology which can be readily deciphered by the receiver, including age (Reby and McComb, 2003; Charlton et al., 2009), sex (Vignal and Kelley, 2007; Charlton et al., 2009), body size (Fitch, 1997; Charlton et al., 2009; Charlton et al., 2011; Garcia et al., 2016), hormone levels (Koren and Geffen, 2009), and physical condition (Wyman et al., 2008; Koren and Geffen, 2009).

In particular, body size often shapes mammalian sounds by constraining the geometry of the vocal tract (Fitch, 2000; Reby and McComb, 2003). Acoustic cues relating to the body size of the caller can inform the receiver about the caller’s competitive ability and reproductive success (Poole, 1999; Reby and McComb, 2003; Kuester et al., 1995; Pfefferle and Fischer, 2006). For example, in primates and carnivores, there is an inverse relationship between body size and call frequency parameters, where larger animals produce calls with lower frequencies, i.e., have a ‘deeper’ voice (Bowling et al., 2017). This relationship between acoustical call features and body size, where one accurately reflects the other, is known as ‘acoustic allometry’ (Taylor and Reby, 2010; Fitch, 1997). Here, signalling is considered *honest* when the acoustic parameters of observed vocalisations accurately reflect an individual’s body size (Zahavi, 1997; Fitch and Hauser, 2003). Deviations from allometry can generate *dishonest* signals, with animals sounding unexpectedly small or large for their body size (Garcia and Ravignani, 2020). Dishonest signals may be produced when an animal shows 1) a lack of allometric scaling between their vocal tract and their body size, or 2) shows enhanced control over their vocal organs which allows them to learn new vocalisations or modify existing vocalisations: an ability known as ‘vocal learning’ (Janik and Slater, 1997; Lattenkamp and Vernes, 2018). Recent work indeed showed that, given a cross-species regression between sounds produced and body size, outlier species seem to cluster either well below the regression line—those with anatomical adaptations—or markedly above—the vocal learners. This led to a morphology vs. learning hypothesis (Garcia and Ravignani, 2020; Ravignani and Garcia, 2021): dishonest signals in mammals may arise either from anatomical adaptations or vocal learning capacities. This prediction has the potential to identify new vocal learners or species with unexpected vocal tract morphology. Vocal learners should therefore be able to violate acoustic allometry while possessing a vocal tract that scales allometrically with the rest of their body. For the first time, we test this prediction, asking whether vocal tract allometry is present in a vocal learning species which is known to violate acoustic allometry.

Harbour seals (*Phoca vitulina*) are vocal learners that escape acoustic allometry by producing sounds with different frequencies than expected from their body size, allowing them to transmit dishonest body size information. Indeed, they stand out as outliers in cross-species allometric regressions between body mass and frequency parameters (Ravignani and Garcia, 2021; see Figure 1). Moreover, previous studies have shown that harbour seals can actively modulate the call frequencies they produce based on auditory experience. In one special case, a human-raised harbour seal, named Hoover, was found capable of mimicking human speech sounds (Ralls et al., 1985). In a more recent study on harbour seal pups, young animals were found capable of lowering their fundamental frequency (f_0_) in the presence of background noise (Torres Borda et al., 2021). Do the environmental noise conditions in which vocalisations are produced have a stronger influence on the f_0_ values than body size? To address this, we complemented acoustic data from Torres Borda and colleagues (2021) with body mass information and reanalysed it to show that acoustic allometric relationships do indeed break down in this species due to the large vocal plasticity observed within individuals (see Figure 2, its caption and detailed explanations in the Supplement). These re-analyses indicate that, also within-species, individual harbour seals may sound bigger or smaller than predicted by body size. Seals can therefore escape the constraints of acoustic allometry, both across and within species.

**Figure 1.**
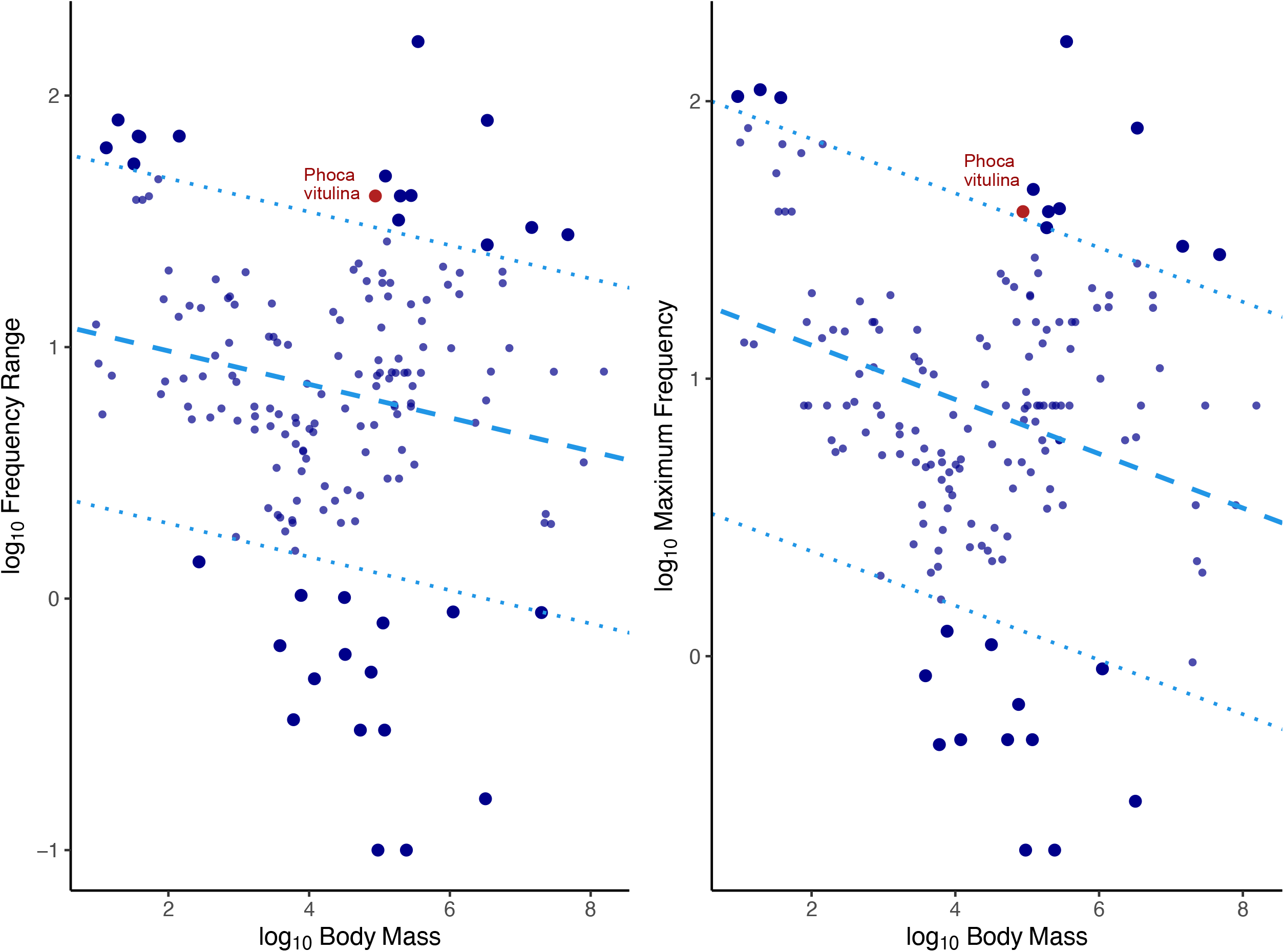
PGLS regressions between frequency parameters (frequency range in the left panel, maximum frequency in the right panel) and body mass across 164 mammalian species. All variables are log-transformed and the figure is adapted from Ravignani and Garcia (2021). The dotted lines represent a threshold at 2,5 standard deviations from the main regression lines used to define outliers. Non-outlier species (which show acoustic allometry between frequency parameters and body mass) are represented by smaller sized circles and outlier species (which escape acoustic allometry) are represented by bigger sized circles. The two red data points, representing harbour seals, are both outliers.

**Figure 2.**
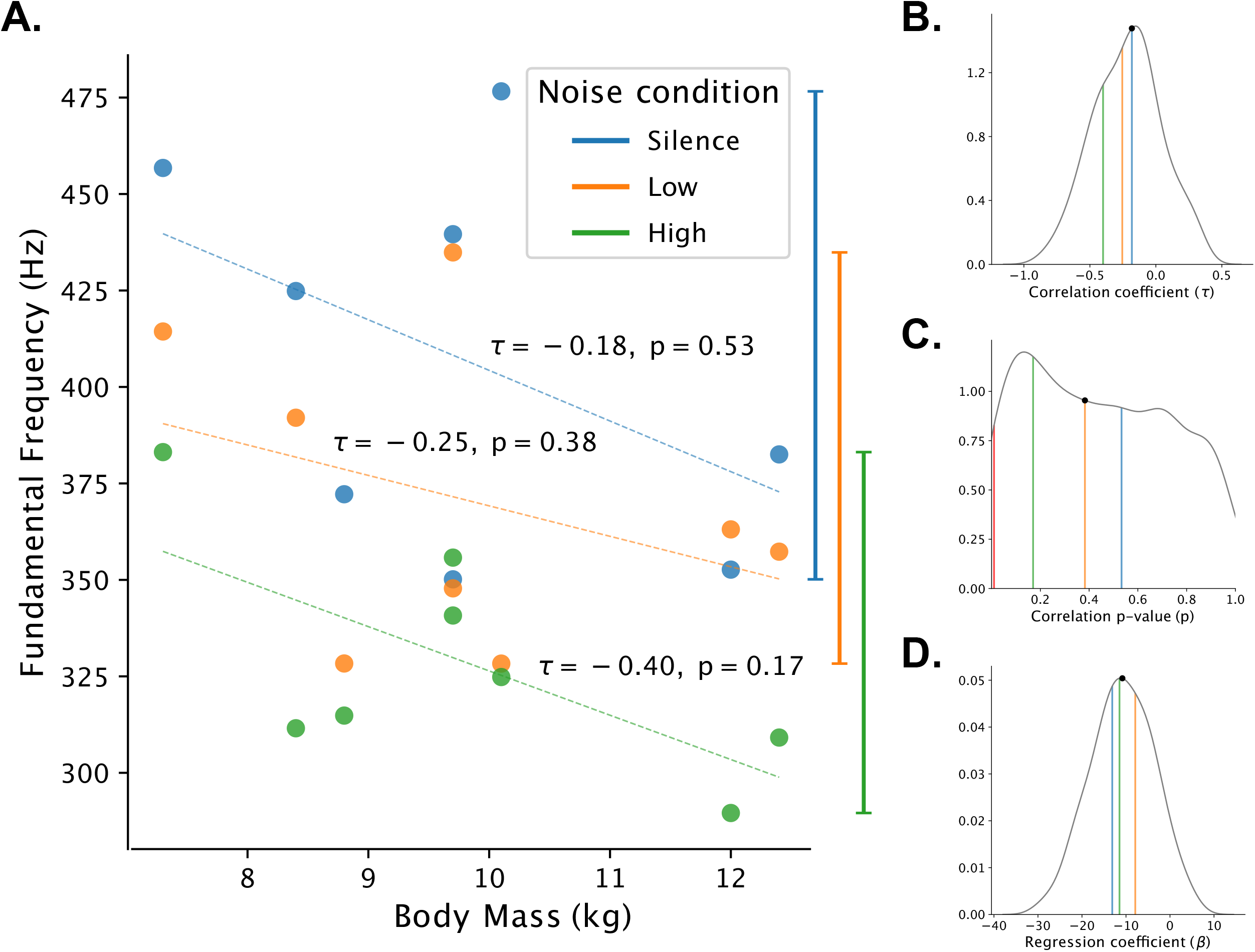
Lack of acoustic allometry relationships in harbour seals. Panel A shows the correlations between median f_0_ for each noise condition (silence, low and high) and body mass. The respective correlation coefficients (τ) and associated p-values (*p*) for each correlation are reported above the regression line. At first sight, the characteristic inverse relationship between f_0_ and body size may seem present, but there is some overlap in the range of f_0_ values (whiskers on the right side of the plot) produced by individuals of differing body size between noise conditions. Non-significant p-values suggest that, at least in this sample, there is a lack of acoustic allometry. In addition, allometry may break if calls are produced in different noise conditions. In other words, do the environmental conditions in which vocalisations are produced strongly affect the f_0_ values, as much as or even more than body mass? We produced density distributions (panels B-D) by computing 10,000 different combinations of randomly selected median f_0_ values (1 of the 3 median frequency values per seal) to assess if allometric relationships hold across noise conditions. The coloured vertical lines in these plots represent the respective values for each of the noise conditions. The median value of the distribution is represented by black circle on the density curve. Panel B shows the density distribution of the Kendall rank correlation coefficients. The median value lies around -0.18, pointing to a weak negative correlation. Panel C shows the density distribution of the correlation p-values associated with the correlations from Panel B. The median p-value is 0.38 which means that in most of the simulated cases we would not reject the null hypothesis (i.e., the correlation is not significantly different from 0). In fact, in only 2.2% of cases (217 out of 10,000) is the correlation significant; this is indicated by the red vertical line. In other words, in 10,000 simulated samples of 8 seals, we generally find no acoustic allometry. Panel D shows the density distribution of the simulated linear regression coefficients, where the median value is -10.8 Hz. Given a 5.1 kg difference in body mass between the smallest and the largest seal, we would expect, on average, a frequency shift of 55.08 Hz. For every individual, we calculated the difference of the median f_0_ values between the silent and high noise condition; the median range across all individuals is 73.6 Hz. This suggests that the differences caused by individual variability in f_0_ in response to noise conditions are larger than the f_0_ differences expected from body mass differences alone. Seals of differing body sizes (e.g., 7 vs. 12 kg) could thus potentially produce the same f_0_ value. This would mean that, in harbour seal pups, vocal plasticity can outweigh and mask acoustic allometric relationships.

Harbour seals are particularly vocal during the first few weeks following birth (Perry and Renouf, 1988). Pups produce individually distinctive mother attraction calls (Renouf, 1984) which vary with age, sex, and body length (Khan et al., 2006; Sauvé et al., 2015). After weaning, however, these calls disappear entirely from their vocal repertoire, with most vocalisations ceasing aside from occasional clicks and growls (Renouf, 1984). During adulthood, female harbour seals remain almost entirely vocally inactive (Van Parijs and Kovacs, 2002), but males start vocalising again, producing underwater calls during the mating season (Hanggi and Schusterman, 1994). The large variation in vocal repertoire observed across individuals, sexes, and age classes makes harbour seals ideal candidates to test the morphology vs. learning hypothesis, i.e., whether a vocal learning mammal does indeed escape acoustic allometry via learning instead of via anatomical adaptations.

Most mammalian vocalizations are described using the source-filter theory of vocal production. Within this framework, vocal signals are initially produced by a *source* and are then *filtered* by the vocal tract before being released into the environment (Fant, 1970). In mammals, the source of sound production consists of the vocal folds in the larynx, and the filter is composed of the cavities making up the upper vocal tract (Fant, 1970) (see Figure 3). The vocal folds are shelves of tissue lying across the airway that attach ventrally and laterally to the thyroid cartilage and dorsally to the arytenoid cartilage (see Figure 4A). When vocalising, the air expelled from the lungs rushes between the vocal folds, causing them to vibrate and produce sound (Elemans et al., 2015). The sound then continues to propagate along the upper vocal tract and is modified by its geometry (i.e., filtered) before being emitted as vocalisation. The source-filter framework highlights which vocal tract structures determine specific features present in acoustic signals. The rate of vibration of the vocal folds determines the f_0_ and the cavities of the vocal tract determine formant frequencies (Taylor and Reby, 2010). Measurements of these vocal tract structures can thus be used to estimate certain acoustic features of vocalisations.

**Figure 3.**
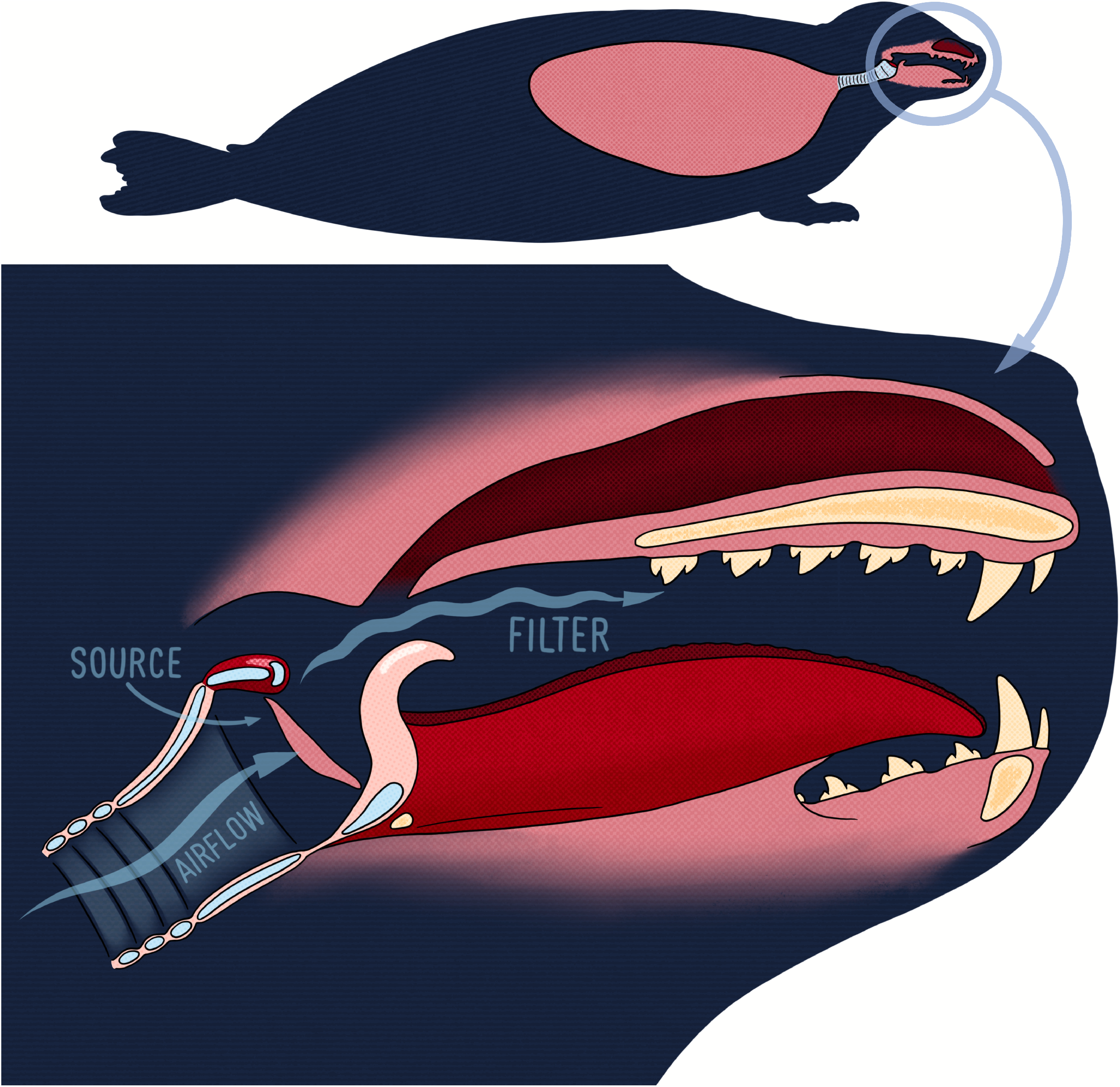
Illustration of the source-filter theory of sound production using the vocal anatomy of the harbour seal.

**Figure 4.**
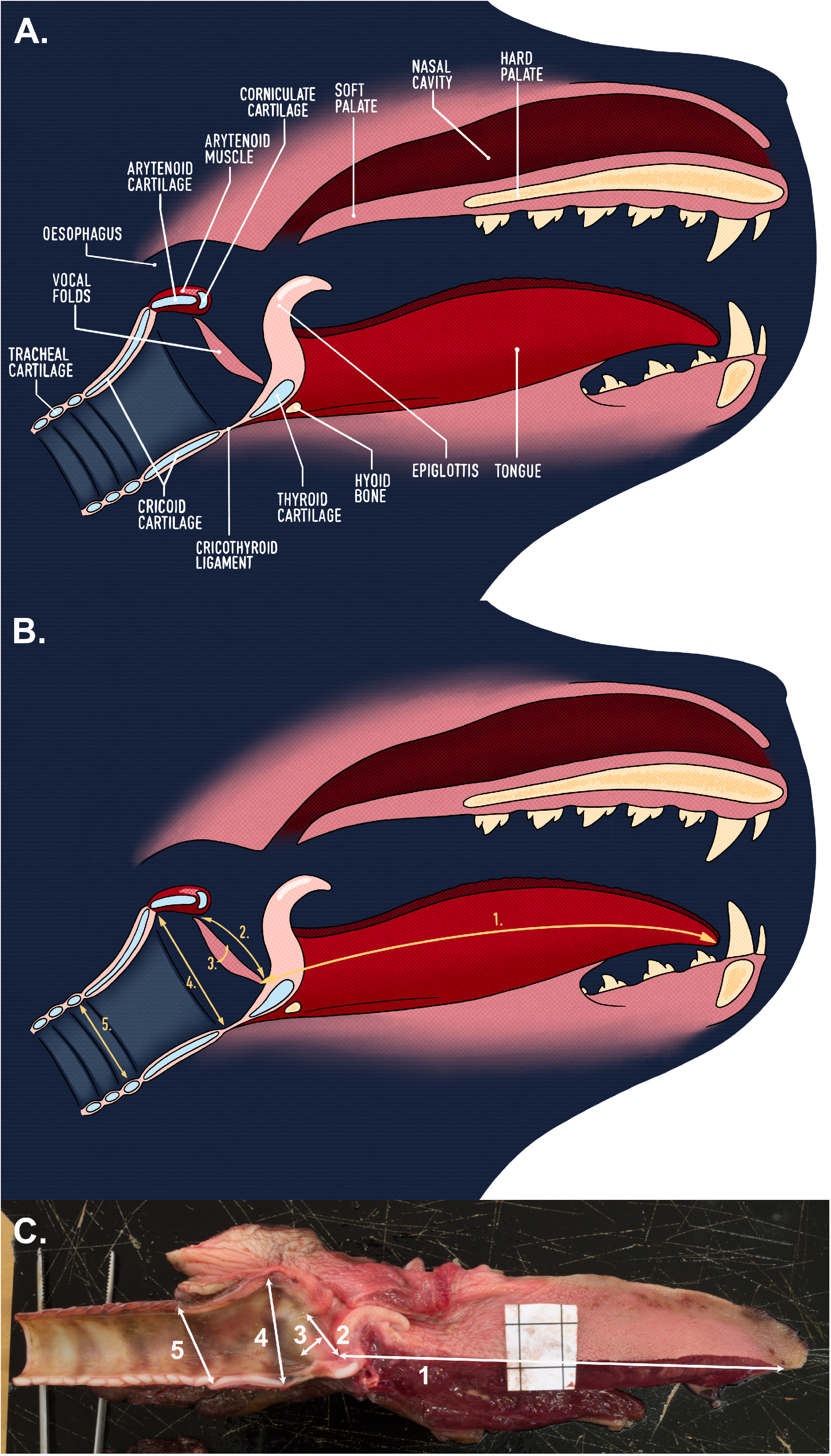
Vocal anatomy of the harbour seal. (A) shows the main anatomical structures composing the vocal tract, (B) depicts the measurements shown on a digital rendering and (C) depicts the measurements shown on a picture of a hemi-larynx from a harbour seal pup. In panel C, the black square outlined on the white paper serves as reference and is exactly 1 cm^2^. The vocal tract measurements taken include (1) vocal tract length (VTL), (2) vocal fold length (VFL), (3) vocal fold thickness (VFT), (4) subglottic-tracheal dorsoventral distance 1 (STDV1), and (5) subglottic-tracheal dorsoventral distance 2 (STDV2).

Bioacoustics studies often investigate allometric relationships between acoustic signal features and body size, without consideration of the underlying allometric scaling between body size and vocal anatomy. Most mammals show allometry between body size and upper vocal tract length because the upper vocal tract is constrained by bony structures (Fitch, 1997; Fitch and Giedd, 1999; Fitch, 2000; Plotsky et al., 2013; Garcia et al., 2016). However, allometry between body size and the size of the vocal folds is less common: the larynx is surrounded by cartilaginous structures and is thus less constrained, suggesting that vocal fold length can be decoupled from overall body size, as found in nonhuman primates (Fitch and Hauser, 1995; Fitch, 1997; Garcia et al., 2017). In mammals, formants, the acoustic proxy of vocal tract length, are thus often a stronger body size predictor than f_0_, the acoustic proxy of vocal fold length (Fitch, 1997; Garcia et al., 2016).

Within the larger framework of the hypothesis above, this study tests for allometric relationships between body size and vocal anatomy measurements in young harbour seals and tests how these relationships vary with sex and age. Preliminary work found that harbour seals’ body length correlates with upper vocal tract length and tracheal diameter, but not with vocal fold length (Ravignani et al., 2017). Here, we aim to expand on these findings by using a larger sample size (353% increase), adding refined anatomical measurements, and comparing different age classes (to test for developmental effects). Based on previous literature, we expect to find allometry between body size and vocal tract structures that are surrounded—and hence constrained—by bony structures, such as vocal tract length. However, based on harbour seals’ vocal learning abilities (Ralls et al., 1985; Torres Borda et al., 2021; Janik and Slater, 1997), we expect their vocal flexibility to offer favourable grounds to find deviations from body size allometry for vocal tract components surrounded by cartilage, such as the trachea and vocal fold length.

## Materials and methods

### Sample collection

Larynges were collected during necropsies on 68 young harbour seals (35 males). Fifty-two samples came from seals that stranded on the Dutch coastline, the rest from animals found on the German coastline (Schleswig-Holstein). Forty-two animals died in captivity at Sealcentre Pieterburen, Pieterburen, the Netherlands, either naturally during rehabilitation despite intensive care or by means of euthanasia due to the presence of severe clinical signs without any indication for recovery. Euthanasia was performed by trained veterinarians, after sedation, with pentobarbital sodium (100 mg/kg) using the method described in Greer and colleagues (2001).

The other 26 animals died in the wild, either naturally or were mercy killed by trained hunters due to severe signs of illness (see Table 1 of the Supplement). No animals were euthanised or mercy killed for the purpose of this study.

**Table 1.** Pairwise Spearman correlations

| Variable | Body Length (cm) | Body Mass (kg) | Girth (cm) | VTL (mm) | VFL (mm) | VFT (mm) | STDV1 (mm) |
| --- | --- | --- | --- | --- | --- | --- | --- |
| Body Mass (kg) | 0.70 |  |  |  |  |  |  |
| Girth (cm) | 0.53 | 0.86 |  |  |  |  |  |
| VTL (mm) | 0.62 | 0.69 | 0.63 |  |  |  |  |
| VFL (mm) | 0.73 | 0.79 | 0.65 | 0.72 |  |  |  |
| VFT (mm) | 0.44 | 0.67 | 0.68 | 0.52 | 0.60 |  |  |
| STDV1 (mm) | 0.63 | 0.78 | 0.65 | 0.70 | 0.82 | 0.69 |  |
| STDV2 (mm) | 0.58 | 0.72 | 0.62 | 0.67 | 0.76 | 0.60 | 0.81 |

*Note.* All correlations were significant at $p < 0.05$ after correcting for multiple comparisons using the Holm-Bonferroni method.

At the time of death, the seals studied were aged between 9 days and 12 months (median 6 months). The age of new-born individuals was estimated in number of days by expert seal veterinarians based on the condition of the umbilical cord or the umbilicus. Older individuals with a closed umbilicus were assigned June as their birth month, which is consistent with the majority of harbour seal births in the Wadden Sea (Osinga et al., 2012; Reijnders et al., 2010). Animals aged 1 month or younger were classified as pups, while those between 1 and 12 months in age were classified as weaners, making age a binary variable. Of the 68 individuals included in this study, 14 (8 males) were classified as pups and 54 (26 males) were classified as weaners. A Fisher’s exact test showed no significant association between age and sex (χ2 = 0.765, *p* > .05), suggesting our sample is balanced between sexes and ages.

### Sample treatment and measurements

Post-mortem examinations were performed by veterinarians who all trained at Sealcentre Pieterburen and thereby used the same necropsy protocol (Pugliares et al., 2007). Dutch seals were examined at Sealcentre Pieterburen and German seals were necropsied at the Institute for Terrestrial and Aquatic Wildlife Research (ITAW), Büsum, Germany. Necropsies were performed on either cooled or defrosted carcasses. Body mass, body length and axillary girth were all measured prior to the start of the necropsy. Body length was measured from the tip of the nose to the end of the tail in a non-curvilinear fashion, while the animal was in supine position, and axillary girth was measured as the body circumference directly caudal to the front flippers. The vocal apparatus including the upper vocal tract, the larynx, and part of the trachea was then removed and immediately frozen at -20°C. All samples were in a similar condition (i.e., none presented signs of decomposition), comparable to pinniped vocal tracts in Schneider (1962) and Ravignani and colleagues (2017).

Prior to measurement, samples were thawed in a refrigerator at 8°C and each larynx was cut medially to produce two hemi-vocal tracts. The measurements taken on these hemi-vocal tracts (see Figures 4B and 4C) include vocal tract length (VTL), vocal fold length (VFL), vocal fold thickness (VFT), and tracheal measurements in the form of subglottic-tracheal dorsoventral distances (STDVs) (called subglottic-tracheal anterior-posterior distance, STAP, in Roers et al., 2009) using a calliper to an accuracy of ±0.01 mm. Although the vocal tract can be divided into lower (below larynx) and upper (above larynx) sections, formants (the resonant frequencies which often encode information about body size) are only determined by the upper vocal tract (Lester and LaGasse, 2008). VTL will henceforth refer to the length of the upper vocal tract. VTL was measured as the linear distance from the caudal end of the epiglottis to the rostral end of the tongue muscle while the tongue was kept straight. VFL was measured as the distance from the ventral attachment of the vocal fold on the thyroid cartilage to the dorsal attachment of the vocal fold on the arytenoid cartilage. VFT was measured as the distance between the anterior and posterior sides of the vocal folds. The first STDV was measured as the distance between the cricothyroid ligament and the caudal end of the arytenoid. The second STDV was measured as the diameter of the first tracheal ring. All measurements were performed independently by two raters (KdR and AR), different from the veterinarians who performed the dissections. For both raters, VTL, VFL, and VFT were measured 4 times, twice for each hemi-vocal tract, and STDVs were taken twice, once for each hemi-larynx, because the start and end measuring points were composed of cartilage (as opposed to soft tissue) and hence, we assumed that the inter-rater reliability for STDVs would be higher than for other measurements.

### Statistical analysis

Statistical analyses were performed in RStudio version 1.1.463 (R version 4.0.4). First, for both raters, the medians for VTL, VFL and VFT were computed from all values reported for every right and left hemi-larynx. Second, using the medians from the first step, the median values for all measurements including STDV1 and STDV2 were computed for each larynx. This provided, for each larynx and rater, five measurements: VTL, VFL, VFT, STDV1 and STDV2. The inter-rater reliability for VTL, VFL, VFT, STDV1 and STDV2 was evaluated using Pearson’s correlations. Finally, the overall median values between raters were computed for all measurements. Using these new values, Spearman’s correlations between body size and vocal anatomy measurements were then calculated (see Table 1). For each measurement, normality was assessed using the Shapiro-Wilk test and homogeneity of variance was assessed using an F-test. If both assumptions were met, a two-tailed independent samples t-test was computed to check for age and sex differences. When variables were not normally distributed, but samples had equal variance, a Mann Whitney U-test was performed to assess group differences instead.

Predictive modelling was done using generalised linear models (GLMs) with the *stats* package (R Core Team, 2013). A series of models were produced for all anatomical measurements with high inter-rater reliability (*r* > 0.70; Salkind, 2010, p. 627). For every response variable, the full model included the fixed effects body length, body mass, girth, sex, age and the interaction effects of sex with all body size predictors, age with all body size predictors and the interaction of age and sex. The reduced model was then obtained through stepwise regression based on Akaike Information Criterion (AIC) values. An analysis of variance (ANOVA) test was performed to ensure that the reduced model was not performing significantly worse than the full one. Variance inflation factors (VIF) scores were calculated for all predictors included in the reduced models using the *car* package (Fox and Weisberg, 2019). Multicollinearity was considered problematic for subsequent model selection if VIF scores were greater than 5 (Akinwande et al., 2015). For all selected models, deviance explained was calculated from the model output (1 – residual deviance / null deviance) and expressed as a percentage. Plots displaying the predicted effects of every predictor retained in the final models were produced to assess their relationship with the response variable. Diagnostic residual plots were used to verify the model assumptions. Independence of residuals was tested using a Durbin Watson test (Fox and Weisberg, 2019). Normality of residuals was assessed visually by plotting model fit against the observed data. Homoscedasticity (i.e., constant variance) of residuals was also assessed visually using quantile-quantile plots. Finally, influential data points were assessed by calculating Cook’s distance.

## Results

Inter-rater reliability for VTL, VFL, VFT, and both STDVs was evaluated using Pearson correlations. VTL (*r* = 0.94), VFL (*r* = 0.88), STDV1 (*r* = 0.97) and STDV2 (*r* = 0.93) showed high inter-rater reliability. VFT (*r* = 0.59) showed lower inter-rater reliability and was consequently excluded from further analysis. All correlations were significant at *p* < 0.001.

All Spearman correlations between body size and vocal anatomy measurements showed positive relationships and significance at the 0.05 level (see Table 1). There were high correlations between body mass and body length (*r*_*s*_ = 0.70), and between body mass and girth (*r*_*s*_ = 0.86). Other notable correlations included those between VTL and VFL (*r*_*s*_ = 0.72), VTL and STDV1 (*r*_*s*_ = 0.70), VFL and STDV1 (*r*_*s*_ = 0.82), VFL and STDV2 (*r*_*s*_ = 0.76). Spearman correlations for pups and weaners can be found in Table 2 of the Supplement.

**Table 2.** Means and standard deviations

|  | All |  | Pups |  | Weaners |  |
| --- | --- | --- | --- | --- | --- | --- |
| Variable | Mean | SD | Mean | SD | Mean | SD |
| Body Length (cm) | 88.07 | 8.03 | 79.86 | 4.79 | 90.19 | 7.32 |
| Body Mass (kg) | 14.53 | 3.92 | 9.81 | 1.46 | 15.75 | 3.39 |
| Girth (cm) | 57.55 | 9.62 | 46.54 | 5.66 | 60.41 | 8.30 |
| VTL (mm) | 91.43 | 6.77 | 83.36 | 4.48 | 93.53 | 5.60 |
| VFL (mm) | 10.92 | 1.16 | 9.26 | 0.90 | 11.35 | 0.77 |
| VFT (mm) | 5.15 | 0.61 | 4.50 | 0.43 | 5.35 | 0.51 |
| STDV1 (mm) | 23.97 | 1.95 | 21.35 | 1.37 | 24.65 | 1.43 |
| STDV2 (mm) | 17.43 | 1.82 | 15.17 | 1.04 | 18.02 | 1.49 |

All anatomical measurements were non-normally distributed but showed equal variances across age and sex groups. A Mann Whitney U-test was used to test for group differences as only the assumption for homogeneity of variance was satisfied. All anatomical measurements were significantly larger in weaners than in pups (*p* < 0.001; see Table 2 and Figure 5). No significant sex differences were found when considering pups and weaners together (*p* > 0.05). When considering pups alone, both the normality and homoscedasticity assumptions were met. A two-tailed independent samples t-test found significant sex differences for vocal tract length (*t* = - 3.42, *p* < 0.05; see Figure 6A). Male pups (86.0 mm ± 2.9) had a larger mean VTL than females (79.8 mm ± 3.7). When considering weaners alone, variables showed non-normal distribution, but equal variances. A series of Mann Whitney U-tests found that only the first subglottic-tracheal dorsoventral distance was significantly different across sexes (*U* = 218, *p* < 0.05; see Figure 6B). Weaned males (25.1 mm ± 1.5) had a wider mean STDV1 compared to weaned females (24.2 mm ± 1.2).

**Figure 5.**
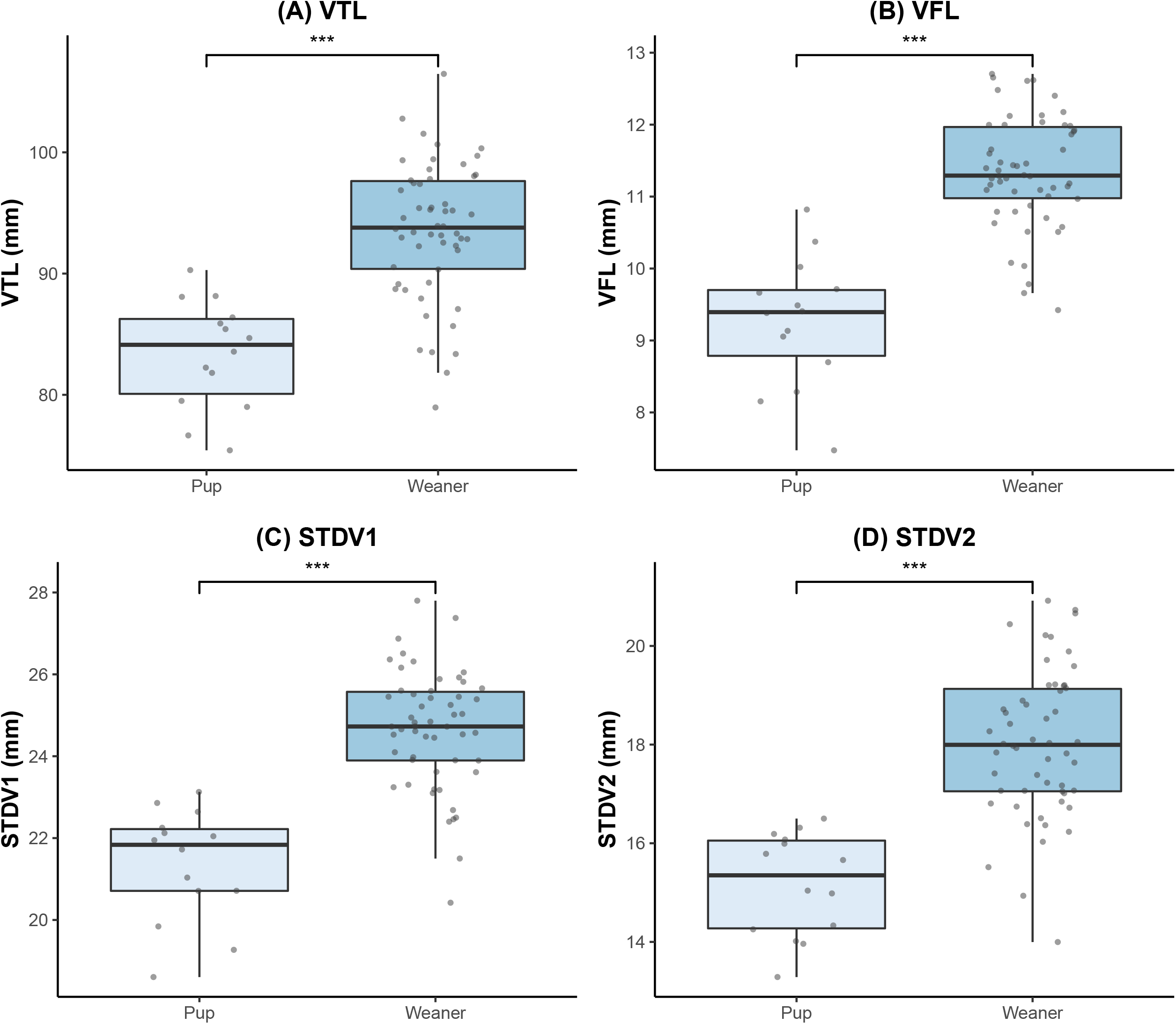
Boxplots illustrating the significant age differences between pups and weaners for (A) VTL, (B) VFL, (C) STDV1, and (D) STDV2. The level of significance is denoted by asterisks, where *** = 0.001.

**Figure 6.**
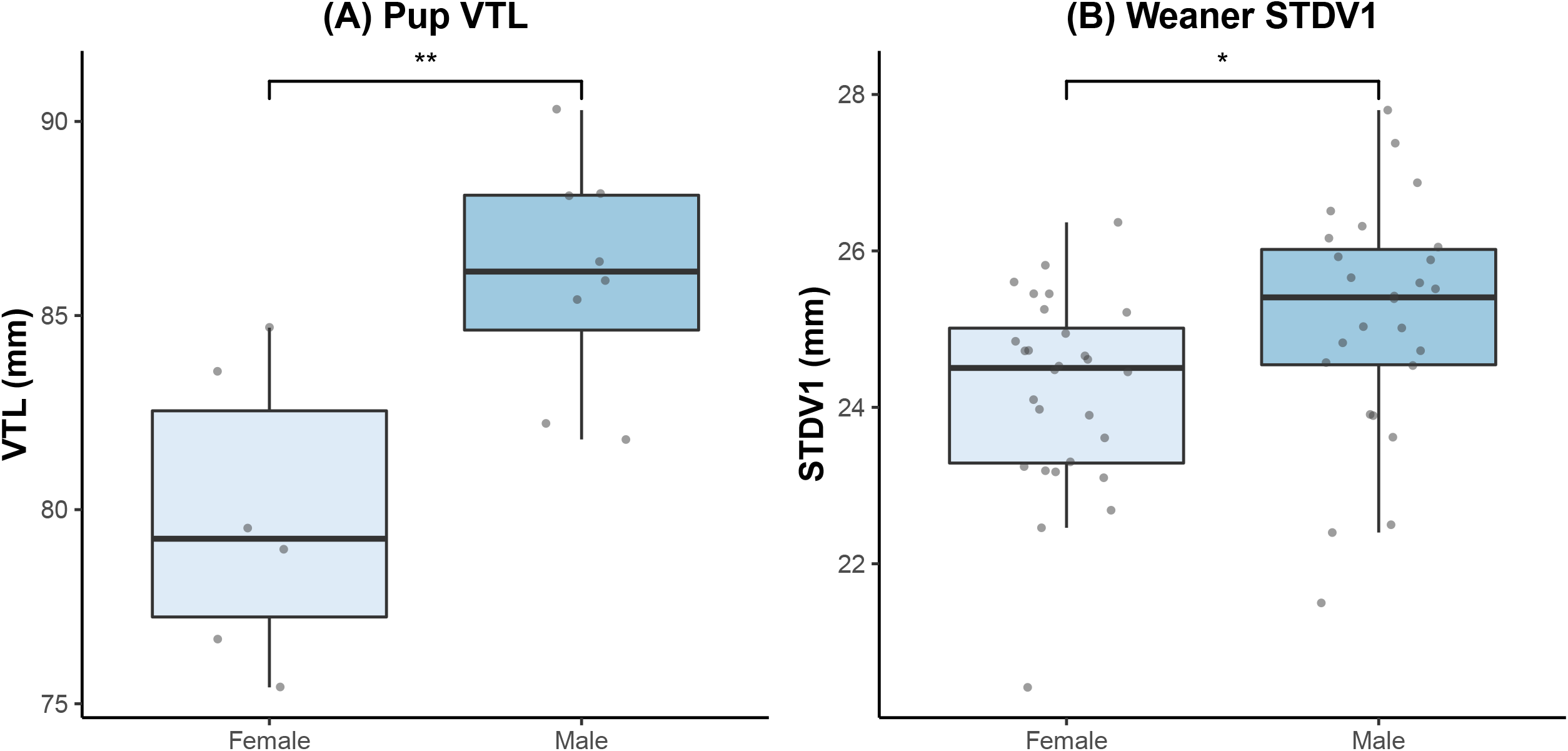
Boxplots illustrating the significant sex differences for (A) VTL in pups and (B) STDV1 in weaners. The level of significance is denoted by asterisks, where * = 0.05 and ** = 0.01.

A reduced GLM, obtained by stepwise regression based on AIC values, was produced for every vocal tract measurement with high inter-rater reliability, including VTL, VFL, STDV1 and STDV2. All VIF scores were lower than 5 suggesting that multicollinearity was not problematic in the selected models. All model assumptions were satisfied. Moreover, ANOVA testing indicated that the reduced models did not perform significantly worse than the full models (*p* > 0.90). GLM results showed that most vocal tract dimensions were best explained by body length, body mass, age, and sex (see Table 3). Girth was not retained as a predictor term in any of the selected models. For each model, the predictor estimates with their confidence intervals can be found in Table 3 of the Supplement and plots of the predicted effects can be found in Figures 1-5 of the same document. Significant interaction effects are shown in Figure 7.

**Table 3.**
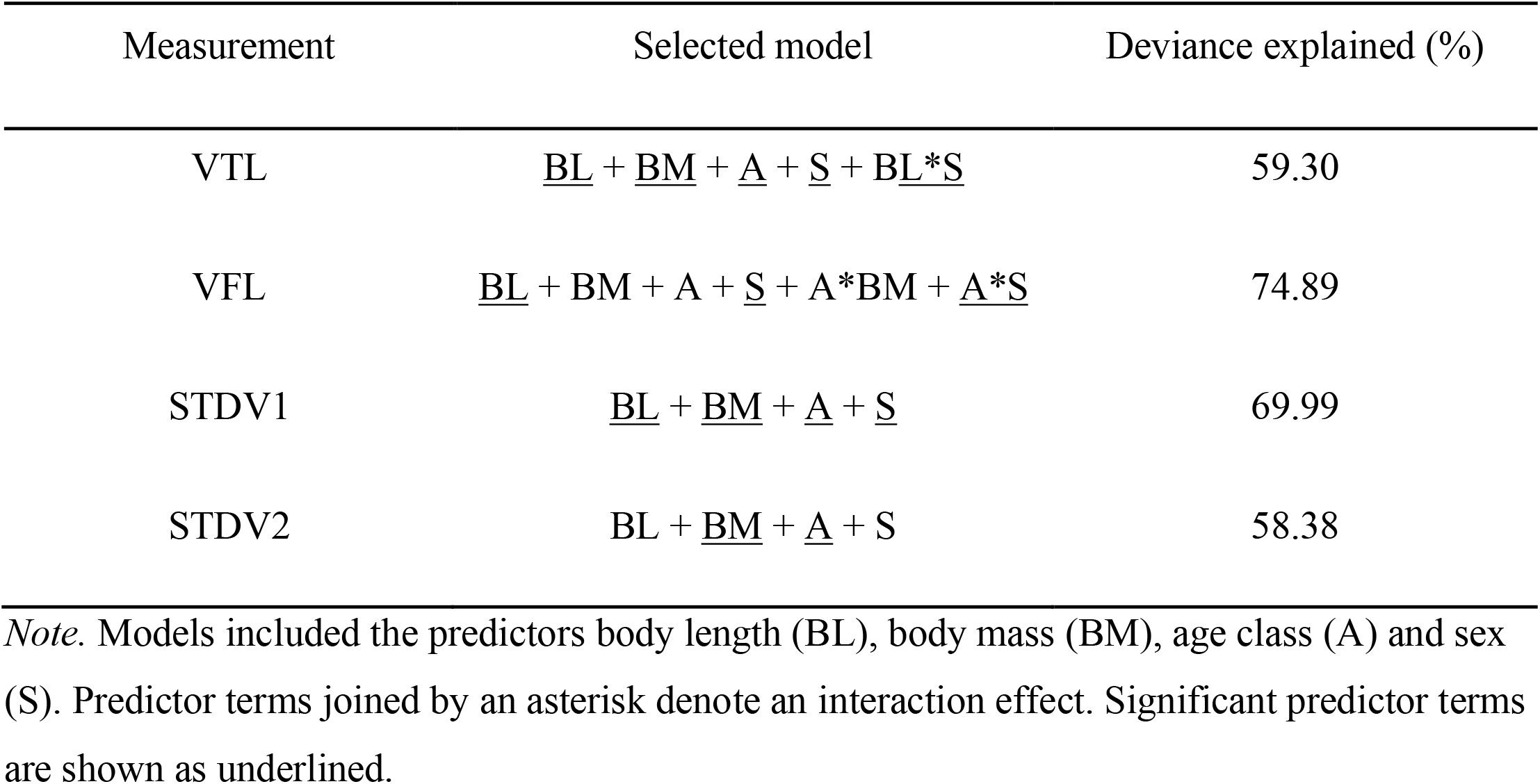
Selected models for each vocal tract structure

**Figure 7.**
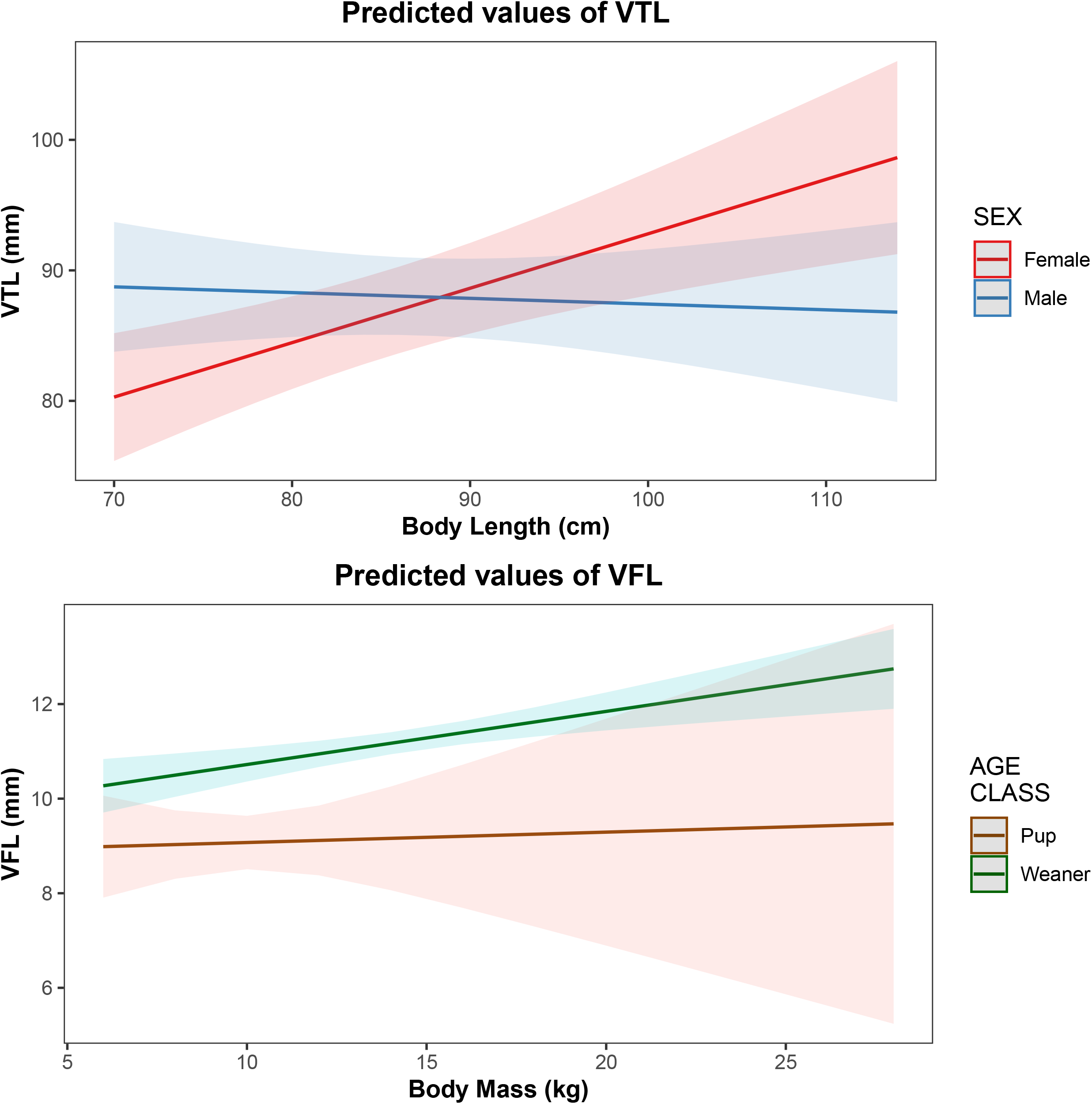
Predicted effects of the body length and sex interaction for VTL including both sexes (top), and the body mass and age interaction for VFL including both age classes (bottom). The shading around each line of best fit indicates the 95% confidence interval.

## Discussion

This study reports on the allometric relationships between body size and vocal tract dimensions in harbour seals. It shows that body length accurately predicts VTL, VFL, and STDV1, and body mass predicts VTL and both tracheal measurements (STDVs). We also find age and sex to be important predictors for the size of vocal tract structures. This is evidenced by significant differences in measurements between age classes and significant sexual differences within age classes.

Previous work showed that upper vocal tract (i.e., filter) dimensions in mammals are predicted by body size measurements (Fitch, 1997; Fitch and Giedd, 1999; Fitch, 2000; Plotsky et al., 2013; Garcia et al., 2016, Ravignani et al., 2017) and our results provide additional evidence to support such allometry. Although most studies have used body length as a proxy for body size, we find that body mass can also be used to predict VTL in harbour seals. In the first years of life, harbour seals show a linear growth rate for both body length (Haukkson, 2006) and mass (Markussen et al., 1989), suggesting that VTL may develop in a similar fashion during this period. Acoustic proxies for the filter could thus provide a good estimation of a harbour seal’s size. In mammals, formant frequencies and formant spacing can be predicted from VTL and vice versa (see Reby and McComb, 2003). Other acoustic proxies include energy quartiles, the frequency of amplitude peaks, and the ratios between these amplitudes (Sauvé et al., 2015). These parameters also encode individual signatures, suggesting that acoustic individuality may partially be an allometric by-product (Ravignani et al., 2017). Harbour seals have the vocal tract predispositions to produce vocalisations that accurately reflect body size whilst also sharing individual-specific information, suggesting that learning does not need to be invoked to explain individuality.

Across mammals, source-related features such as f_0_ can sometimes predict body size despite showing weaker allometric scaling than filter-related features (Reby and McComb, 2003; Charlton et al., 2011; Pfefferle et al., 2007; Charlton and Reby, 2016); it was unclear whether this holds for harbour seals (Ravignani et al., 2017; Bowling et al., 2017). Our findings indicate that VFL, which may be used to approximate f_0_, can be predicted by body size in harbour seals. Moreover, Sauvé and colleagues (2015) reported a decrease in f_0_ with an increase in body length of harbour seal pups. Taken together, this suggests that a harbour seal’s f_0_ can be predicted from vocal anatomy. Previous evidence against allometric scaling for VFL could be explained by low statistical power or lack of testing for age effects on vocal tract measurements (Ravignani et al., 2017). It is indeed notable that age is included in both interactions which were retained in the selected VFL model. Our results, including both pups and weaners, show that allometric scaling between body size and VFL only emerges after weaning, suggesting that VFL may not be constrained in harbour seal pups (see bottom panel of Figure 7). This begs the following question: how would escaping acoustic allometry for source-related features be beneficial for pups? Broadcasting honest body size information may be detrimental for harbour seal pups as they are significantly more likely to be displaced by larger conspecifics during agonistic interactions (Neumann, 1999). However, pups may be able to benefit from lowering the f_0_ (Torres Borda et al., 2021) of their calls to create an impression of size exaggeration. On the other hand, pups may also benefit from increasing the f_0_ of their calls to create an impression of distress to the mother (Briefer, 2012). Future playback studies could and should contrast these hypotheses.

Several phocid species use the trachea for sound production (Bryden and Felts, 1974), but this could be a by-product of adaptive modifications to the respiratory tract required for diving (Kooyman and Andersen, 1969; Tyack and Miller, 2002). Our results support the correlation between tracheal diameter and body length found by Ravignani and colleagues (2017), but also provide evidence that tracheal dimensions can be predicted by body mass. Previous literature found that the trachea may potentially convey body size information if its size influences acoustic call features (Ravignani et al., 2017). In humans, a wider tracheal diameter partially predicts turbulence (i.e., unsteady air movements) for large airflows (Van den Berg et al., 1957). Applying the same logic to other mammals, larger seals would have wider tracheal dimensions which, in turn, would make vocalisations noisier. This could explain, for instance, why the harmonics-to-noise ratio decreases as harbour seals get older (de Reus, 2017). Future work on sound production in this species could test this prediction using sound-anatomy correlations and excised larynx set-ups. Moreover, playback experiments could test whether adding noise to vocalisations alters interactive behaviour to determine if harmonics-to-noise ratio may encode body size information. Understanding whether and how the trachea is involved in sound production will thus require further research.

As expected, all anatomical measurements are larger for weaners than they are for pups. In Ravignani and colleagues (2017), animals up to 108 days old were classified as pups. However, in the wild, the lactation period for harbour seals ranges from 23 to 42 days, after which the pups are weaned (Renouf, 2012). Hence, for the sake of simplicity, we consider animals up to one month old as pups and animals older than one month as weaners. Through this categorical classification, we were able to identify how allometric trends develop over the harbour seal’s early life. At the time of data collection, we only had very few larynges from subadults and adults, leading us to not include these data points in our analysis to avoid potential problems caused by small sample size. Future research including larynges from subadults and adults will further extend our knowledge of how vocal allometry develops in harbour seals.

There were no sexual differences when considering the sample size as a whole, but significant sexual differences existed within age classes. These differences could be attributed to differing levels of steroid hormones acting on the laryngeal structures in males and females (Aufdemorte et al., 1983; Sauvé et al., 2015). In some mammals, sex hormones affect the structural development of the larynx and the viscoelastic properties of the vocal fold tissue (Fitch and Giedd, 1999; Beckford et al., 1985). At puberty of these animals, the male larynx descends in the vocal tract causing an elongation of the length of the upper vocal tract, allowing males to convey an exaggerated impression of size (Fitch and Giedd, 1999; Fitch and Reby, 2001). In harbour seal pups of similar body size, males have larger VTLs than females, suggesting that laryngeal descent in males possibly occurs early in life. Once weaned, however, females show a clear increase in VTL whereas it remains relatively constant in males (see top panel of Figure 7), suggesting that VTL differences across sexes may become less pronounced over time. In mammalian males, sex hormone action also causes a rapid increase in cartilage size leading to an enlarged larynx and an increase in the vibrating portion of the vocal folds (Fitch and Hauser, 2003). This could explain why, in weaners, STDV1 is larger in males than in females. Nevertheless, these findings are somewhat surprising as young harbour seals normally show little sexual dimorphism (Le Boeuf, 1991). In particular, there is a lack of evidence for sexual differences regarding birth mass and growth rates among harbour seal pups (Bowen et al., 1994). In our sample, there are no significant body size differences between sexes *(p* > 0.05), however, male pups are slightly larger than female pups in body length (M = 81.6 cm ± 4.4, F = 77.5 cm ± 4.5), which could partially explain the VTL differences observed in this age class. Male (9.8 kg ± 1.5) and female (9.8 kg ± 1.6) pups do not differ in body mass, but it is important to note that the sampled animals were sick and/or in poor condition; hence body mass values are not representative of healthy individuals and should be interpreted with caution. In short, based on these observed differences in vocal anatomy across sexes, formants are expected to differ in pups and harmonics-to-noise ratio is expected to differ in weaners. The anatomical structures that determine these acoustic features both show strong allometric scaling, hence these parameters may provide distinct body size cues across age classes, potentially facilitating the discrimination of male and female conspecific calls. Future research should investigate how sex hormones affect the elastic properties of harbour seal laryngeal tissues. Hormone levels can be measured by taking blood samples from healthy male and female seals at different developmental stages, and results can be combined with magnetic resonance imaging (MRI) mapping of laryngeal tissue elasticity.

The high inter-rater reliability observed for VTL, VFL and both STDVs demonstrates that these quantities can be measured and replicated easily, making them reliable landmarks for vocal tract measurements. However, tissue properties such as the viscoelasticity of certain vocal tract structures, like the vocal folds, are significant obstacles to getting accurate measurements. Indeed, raters struggled to produce precise data for VFT. Future research in the field of pinniped vocal anatomy would benefit from improved measuring techniques using 2D pictures, radiography, MRI and computed tomography scans as this would enable more accurate measurements for structures that are difficult to handle. Finally, future similar studies should include measurements of another vocal tract structure: the corniculate cartilage. Although widely absent in terrestrial carnivores, harbour seals have rather large corniculate cartilages that help close the trachea together with the epiglottis (Adams et al., 2020). These cartilages are located close to the vocal folds and are possibly innervated by the same nerves and controlled by the same muscles. It may be possible that these cartilages play a role in sound production by, for example, lowering the f_0_ by adding weight to the vocal folds. Taken together, these suggestions will provide a more precise and detailed picture of the harbour seal’s vocal anatomy.

Observed species-specific vocalisations are determined by both the species’ vocal anatomy and their capacity for vocal learning (Garcia and Ravignani 2020; Ravignani and Garcia 2021). The vocal anatomy generates vocal predispositions by imposing biomechanical constraints, whereas neural processes determine the degree of control species have over their vocal organs (Garcia and Manser, 2020). Particularly, vocal learners, like the harbour seal, are capable of actively modulating sounds, suggesting that they are less constrained by anatomy and have a refined capacity for vocal motor control. Unfortunately, the relative contribution of both sound production mechanisms is unclear. Here, we test a hypothesis trying to segregate anatomical vs. learning mechanisms (Garcia and Ravignani, 2020; Ravignani and Garcia 2021). As shown here, by testing for allometric relationships between body size and vocal tract structures, one can start to disentangle the respective contributions of vocal anatomy and vocal motor control in shaping acoustic signals. We find that harbour seals are mechanistically constrained by their vocal anatomy, and their large vocal flexibility (Ralls et al., 1985; Torres Borda et al., 2021), which may result in the production of dishonest signals, thus points towards extensive volitional control over their vocalisations. In brief, we provide support for the morphology vs. learning hypothesis, showing however that this relation may be mediated by ontogeny.

In sum, we provide evidence of allometry between body size and vocal tract measurements in harbour seals. Body length is a strong predictor for VTL, VFL, and STDV1, and body mass is a strong predictor for VTL and both tracheal measurements (STDVs). Age and sex are also important in predicting the dimensions of these anatomical structures. Taken together, the combined findings demonstrate that harbour seal vocal tracts do indeed scale with body size, although allometry between VTL and body size may only emerge after weaning. One could now make inferences about the vocal predispositions of harbour seals (e.g., f_0_, formants), based on either their body size or the size of their vocal tract. However, to accurately predict f_0_, further studies are needed in harbour seals to determine the range of stress they apply to their vocal folds while vocalising and to infer the tissue density of their vocal folds (Titze et al., 1989). Once such predictions are made, comparing them to data obtained from observed natural vocalisations would shed light on the range of vocal flexibility resulting from their extensive vocal motor control. Although formant spacing could be predicted from vocal tract length (Titze, 1994), bioacousticians have not yet been able to consistently extract formants from harbour seal vocalisations, meaning that predictions cannot currently be compared to observed vocalisations. Finally, a critical next step to directly relate acoustic features to sound production structures is to connect harbour seals’ vocal anatomy measurements to the vocalisations they produce while alive. Integrating such results with investigations of call function will eventually inform on which vocal structures are responsible for generating the individual- and species-specific information encoded in harbour seals’ vocalisations.

## List of symbols and abbreviations

f_0_: Fundamental frequency
C: Celsius
VTL: Vocal Tract Length
VFL: Vocal Fold Length
VFT: Vocal Fold Thickness
STDV: Subglottic-Tracheal Dorsoventral distance
STAP: Subglottic-Tracheal Anterior-Posterior distance
GLM: Generalised Linear Model
AIC: Akaike Information Criterion
ANOVA: Analysis Of Variance
VIF: Variance Inflation Factor
M: Male
F: Female
mg: milligram
kg: kilogram
cm: centimetre
MRI: Magnetic Resonance Imaging

## Acknowledgments

We are grateful to Letty Stupers, Rebecca Andreini, Maria Jose Robles for help in gathering and preparing samples for data collection. We thank Aline Joustra for her illustrations of seal vocal tracts. We also greatly appreciate the extensive comments provided by Taylor Hersh, Laura Verga and Yannick Jadoul. We would also like to thank all members and volunteers of Sealcentre Pieterburen for their continued support in this research.

## Author contributions

KdR, SG, MG, ARG, ASC and AR conceived the research and designed the experiment. SG, ARG, and ASC performed the post-mortem examinations. KdR and AR performed the measurements. KdR, DC, and AL performed the statistical analysis. All authors interpreted the results, drafted the article, and helped to critically revise it. KdR and AR addressed the reviewers’ comments and incorporated them in the manuscript with support from SG, MG, ARG, and ASC.

