## Supplement for "Vocal tract allometry in a mammalian vocal learner"

### Acoustic analyses

We ran several acoustic analyses to show that seals can escape acoustic allometry. The acoustic data used to perform these analyses comes from the study published by Torres Borda and colleagues (2021), during which they observed fundamental frequency changes in harbour seal pups under different noise conditions (silence, low and high noise). The acoustic data was complemented with previously unpublished body mass measurements of the animals in their study. The harbour seal pups were weighed on their day of arrival at the Sealcentre Pieterburen, a pinniped rehabilitation centre in the Netherlands, where the animals were also audio recorded. Eight harbour seals participated in the noise playback experiment and the fundamental frequency ($f_0$) was extracted from the vocalisations they produced during the testing period (all details in Torres Borda et al., 2021).

Grouping the observations by seal ID and noise condition, we computed the median $f_0$ for each of the 24 groups. We then regressed median $f_0$ on body size (using body mass as a proxy for body size) for each of the noise conditions (Figure 2A). Visually, an inverse relationship between body size and call frequency seems to hold in all three noise conditions, but none of the correlations ($\tau_{silence} = -0.18$, $\tau_{low} = -0.25$, $\tau_{high} = -0.40$) are significant ($p < 0.05$). This apparent inconsistency may be explained by a large degree of overlap in the range of $f_0$ values produced by individuals of differing body size between noise conditions. For instance, an animal of 12.4 kg under silence can produce a similar $f_0$ value as an animal of 7.3 kg under high noise (see Figure 2A), suggesting that acoustic allometry may not hold across noise conditions. Could it be that the environmental noise conditions in which vocalisations are produced more strongly affect the $f_0$ values than body size? If hypothetically we were to record calls of harbour seal pups on different days and irrespective of environmental noise conditions, the inverse relationship between $f_0$ and body size may disappear (i.e., acoustic allometry would break) if the individuals can, thanks to their large vocal plasticity, adjust their $f_0$ depending on the noise conditions.

To assess if allometric relationships do indeed break down across noise conditions, we computed 10,000 different combinations of randomly selected median $f_0$ values (1 of the 3 median frequency values per seal) and matched each value to the corresponding body mass value. We then performed 10,000 Kendall rank correlations, each among the 8 resulting pairs of $f_0$ and body mass values. Figure

2B shows the kernel density distribution of the resulting correlation coefficients and their associated p-values (Figure 2C). We find that the median correlation coefficient is -0.18, suggesting a weak negative correlation. The median p-value is 0.38, indicating that—in more than half of the cases—we cannot reject the null hypothesis which states that the correlations are generally not significantly different from 0. In only 2.2% of cases (217 out of 10,000) is the correlation significant. We should take care when interpreting the correlation p-values as the power of the test statistic is low given the small sample size (n = 8), resulting in a higher probability of committing type II errors. Moreover, the body mass values correspond to the measurements taken on the day of the animal's arrival at the Sealcentre; they are not representative of the actual body mass values on the days of testing. Using the same set of random combinations of $f_0$ values, we also plotted the density distribution for the linear regression coefficients (Figure 2D). The median regression coefficient is -10.8 Hz/kg. The difference in initial body mass between the largest and smallest seal is 5.1 kg. This means that across their mass range, we would expect, on average, a 55.08 Hz difference. For every seal, we calculated the range between the median $f_0$ values of the silent and high noise condition (silence $f_0$ – high $f_0$) and find that the median is 73.6 Hz. This suggests that the differences caused by individual variability in $f_0$ in response to noise conditions are larger than the $f_0$ differences expected from body mass differences alone. Seals of differing body sizes (e.g., 7 vs. 12 kg) could thus potentially produce the same $f_0$ value (and they actually do, see Figure 2A). Furthermore, we also calculated, for each seal, the $f_0$ range (maximum – minimum $f_0$) for all recorded observations from that individual. We find that, across the tested seals, the median $f_0$ range is 322.6 Hz. Applying the same logic as above, seals with a body mass difference of almost 30 kg (322.6 / 10.8) could all produce similar $f_0$ values. Finally, we computed and compared two simple generalised linear models, testing if body mass (Model 1: $f_0$ ~ Body Mass) or noise condition (Model 2: $f_0$ ~ Noise Condition) was better at predicting $f_0$. We find that body mass is not a significant predictor of $f_0$ ($t$ = -1.78, $p$ = 0.09), but noise condition is ($t_{high\ vs.\ low}$ = 2.10, $p$ = 0.048; $t_{high\ vs.\ silence}$ = 3.90, $p$ = 0.001). Moreover, Model 1 explained 12.63% of the deviance (calculated as (1 - residual deviance / null deviance) * 100) and Model 2 explained 42.05% of the deviance. An ANOVA test confirmed that Model 2 significantly outperformed Model 1 (F = 10.7, $p$ = 0.001), showing that environmental noise conditions may have a stronger influence on $f_0$ than body size.

Table 1

*List of sampled animals*

| ID | Age class | Where from | Sex | Body Length (cm) | Body Mass (kg) | Girth (cm) | Cause of death |
|---|---|---|---|---|---|---|---|
| 1 | weaner | NL | F | 86 | 15.6 | 90 | Euthanised |
| 2 | weaner | NL | F | 99 | 17.3 | 83 | Died during rehab |
| 3 | weaner | NL | M | 96 | 26.8 | 76 | Found dead in the wild |
| 4 | weaner | NL | M | 96 | 22.9 | 71 | Euthanised |
| 5 | weaner | NL | M | 86 | 19.2 | 71 | Died before rehab |
| 6 | weaner | NL | M | 84 | 18.2 | 69 | Euthanised |
| 7 | weaner | NL | M | 89 | 19.9 | 66 | Euthanised |
| 8 | weaner | NL | F | 94 | 14.2 | 66 | Euthanised |
| 9 | weaner | NL | F | 92 | 15.8 | 65 | Euthanised |
| 10 | weaner | NL | F | 94 | 14.9 | 65 | Died during rehab |
| 11 | weaner | NL | M | 104 | 20.8 | 64 | Died before rehab |
| 12 | weaner | NL | F | 86 | 16.5 | 63 | Died during rehab |
| 13 | weaner | NL | F | 86 | 15.7 | 63 | Euthanised |
| 14 | weaner | NL | M | 100 | 18.37 | 62 | Euthanised |
| 15 | weaner | NL | F | 93 | 16.8 | 62 | Euthanised |
| 16 | weaner | NL | M | 114 | 18.8 | 61 | Died before rehab |
| 17 | weaner | NL | F | 87 | 15.8 | 61 | Euthanised |
| 18 | weaner | NL | F | 93 | 17.8 | 60.5 | Euthanised |
| 19 | weaner | NL | M | 96 | 16.3 | 60 | Died during rehab |
| 20 | weaner | NL | F | 82 | 15.3 | 60 | Euthanised |
| 21 | weaner | NL | M | 80 | 14.3 | 60 | Died before rehab |
| 22 | weaner | NL | F | 88 | 16.1 | 59 | Died during rehab |
| 23 | weaner | NL | F | 89 | 16.9 | 58 | Euthanised |
| 24 | weaner | NL | F | 71 | 10 | 58 | Euthanised |
| 25 | weaner | NL | M | 92 | 17 | 57 | Euthanised |
| 26 | weaner | NL | F | 94 | 14.5 | 57 | Died during rehab |
| 27 | weaner | NL | M | 86 | 13.9 | 57 | Died before rehab |
| 28 | weaner | NL | F | 79 | 11.9 | 56.5 | Euthanised |
| 29 | weaner | NL | M | 85 | 14.6 | 56 | Died during rehab |
| 30 | weaner | NL | M | 94 | 14.6 | 55 | Euthanised |
| 31 | weaner | NL | F | 92 | 13.7 | 55 | Died during rehab |
| 32 | weaner | NL | M | 80 | 13.1 | 55 | Died during rehab |
| 33 | pup | NL | F | 75 | 11.9 | 55 | Euthanised |
| 34 | pup | NL | M | 84 | 11.79 | 54 | Found dead in the wild |
| 35 | weaner | NL | F | 93 | 14 | 53 | Died during rehab |
| 36 | pup | NL | M | 83 | 11.47 | 52 | Euthanised |
| 37 | weaner | NL | F | 93 | 13.9 | 51.5 | Euthanised |

| | | | | | | | |
|---|---|---|---|---|---|---|---|
| 38 | weaner | NL | M | 86 | 13 | 51.5 | Died before rehab |
| 39 | weaner | NL | F | 87 | 12.4 | 51 | Died before rehab |
| 40 | pup | NL | M | 86 | 10.6 | 51 | Euthanised |
| 41 | pup | NL | F | 81 | 11.37 | 49.5 | Euthanised |
| 42 | pup | NL | F | 82 | 9.3 | 49 | Found dead in the wild |
| 43 | pup | NL | M | 80 | 9.46 | 47 | Euthanised |
| 44 | pup | NL | M | 73 | 8.6 | 46 | Found dead in the wild |
| 45 | pup | NL | F | 77 | 8.5 | 44.5 | Died before rehab |
| 46 | pup | NL | F | 80 | 9.63 | 44 | Found dead in the wild |
| 47 | weaner | NL | M | 87 | 9.3 | 44 | Died during rehab |
| 48 | pup | NL | F | 70 | 8 | 44 | Found dead in the wild |
| 49 | pup | NL | M | 87 | 9.43 | 41 | Found dead in the wild |
| 50 | weaner | NL | F | 77 | 7.47 | 40 | Euthanised |
| 51 | pup | NL | M | 80 | 7.28 | 38.5 | Died during rehab |
| 52 | pup | NL | M | 80 | 9.95 | 36 | Died before rehab |
| 53 | weaner | DE | M | 85.5 | 19.2 | 67.5 | Mercy killed |
| 54 | weaner | DE | F | 98.5 | 17 | 66 | Mercy killed |
| 55 | weaner | DE | M | 90 | 14.6 | 65 | Mercy killed |
| 56 | weaner | DE | M | 101 | 20.8 | 64 | Mercy killed |
| 57 | weaner | DE | M | 90 | 17 | 63 | Found dead in the wild |
| 58 | weaner | DE | F | 92 | 20.4 | 62 | Found dead in the wild |
| 59 | weaner | DE | F | 99 | 17.8 | 60.5 | Mercy killed |
| 60 | weaner | DE | M | 86 | 16 | 60.5 | Found dead in the wild |
| 61 | weaner | DE | M | 90 | 16.6 | 59 | Found dead in the wild |
| 62 | weaner | DE | F | 94 | 17 | 58 | Found dead in the wild |
| 63 | weaner | DE | M | 90 | 14.6 | 57 | Mercy killed |
| 64 | weaner | DE | M | 96 | 13.4 | 56 | Found dead in the wild |
| 65 | weaner | DE | F | 82 | 11.6 | 55.5 | Found dead in the wild |
| 66 | weaner | DE | F | 97 | 14.8 | 54 | Mercy killed |
| 67 | weaner | DE | M | 81 | 10.2 | 51 | Mercy killed |
| 68 | weaner | DE | F | 88.5 | 12 | 50 | Mercy killed |

*Note.* Seals were from the Netherlands (NL) or Germany (DE). Sex is denoted as F for females and M for males.

Table 2

*Pairwise Spearman correlations for pups and weaners*

| Age class | Variable | Body Length (cm) | Body Mass (kg) | Girth (cm) | VTL (mm) | VFL (mm) | VFT (mm) | STDV1 (mm) |
|---|---|---|---|---|---|---|---|---|
| Pups | Body Mass (kg) | 0.40 | | | | | | |
| | Girth (cm) | 0.23 | 0.72 | | | | | |
| | VTL (mm) | 0.23 | -0.03 | -0.04 | | | | |
| | VFL (mm) | 0.22 | 0.08 | -0.01 | 0.49 | | | |
| | VFT (mm) | -0.21 | 0.11 | 0.42 | 0.01 | 0.16 | | |
| | STDV1 (mm) | 0.20 | 0.18 | 0.08 | 0.49 | 0.79* | 0.55 | |
| | STDV2 (mm) | 0.35 | 0.36 | 0.38 | 0.26 | 0.71 | 0.57 | 0.76* |
| Weaners | Body Mass (kg) | 0.51* | | | | | | |
| | Girth (cm) | 0.28 | 0.76* | | | | | |
| | VTL (mm) | 0.39* | 0.48* | 0.39* | | | | |
| | VFL (mm) | 0.58* | 0.64* | 0.41* | 0.54* | | | |
| | VFT (mm) | 0.16 | 0.48* | 0.50* | 0.22 | 0.34 | | |
| | STDV1 (mm) | 0.38* | 0.61* | 0.38* | 0.46* | 0.67* | 0.47* | |
| | STDV2 (mm) | 0.32 | 0.48* | 0.31 | 0.47* | 0.57* | 0.30 | 0.66* |

*Note*. * indicates $p < .05$ after correcting for multiple comparisons using the Holm-Bonferroni method.

Table 3

*Generalised linear model (GLM) estimates for all vocal structures*

| Vocal structure | Effect | Estimate | Std.Err. | 2.5% | 97.5% | p |
|---|---|---|---|---|---|---|
| VTL | Intercept | 42.4788 | 9.2445 | 23.9898 | 60.9678 | < 0.001 |
| | Age Class-Weaner | 4.6695 | 1.7801 | 1.1093 | 8.2297 | < 0.05 |
| | Body Length | 0.4170 | 0.1184 | 0.1802 | 0.6538 | < 0.001 |
| | Body Mass | 0.5933 | 0.2182 | 0.1569 | 1.0297 | < 0.01 |
| | Sex-Male | 40.7192 | 12.1685 | 16.3822 | 65.0562 | < 0.01 |
| | Body Length*Sex-Male | -0.4610 | 0.1379 | -0.7368 | -0.1852 | < 0.01 |
| VFL | Intercept | 9.1651 | 3.0598 | 3.0455 | 15.2847 | < 0.01 |
| | Age Class-Weaner | -2.7492 | 3.2348 | -9.2188 | 3.7204 | 0.399 |
| | Body Length | -0.0189 | 0.0396 | -0.0981 | 0.0603 | 0.635 |
| | Body Mass | 0.1050 | 0.0301 | 0.0448 | 0.1652 | < 0.001 |
| | Sex-Male | 1.0070 | 0.3670 | 0.2730 | 1.7410 | < 0.01 |
| | Age Class-Weaner*Body Length | 0.0562 | 0.0411 | -0.026 | 0.1384 | 0.177 |
| | Age Class-Weaner*Sex-Male | -1.1833 | 0.4071 | -1.9975 | -0.3691 | < 0.01 |
| STDV1 | Intercept | 15.389 | 1.6827 | 12.0236 | 18.7544 | < 0.001 |
| | Age Class-Weaner | 1.7474 | 0.4352 | 0.8770 | 2.6178 | < 0.001 |
| | Body Length | 0.0472 | 0.0236 | 0.0000 | 0.0944 | < 0.05 |
| | Body Mass | 0.1887 | 0.0533 | 0.0821 | 0.2953 | < 0.001 |
| | Sex-Male | 0.5956 | 0.2754 | 0.0448 | 1.1464 | < 0.05 |
| STDV2 | Intercept | 9.9726 | 1.8526 | 6.2674 | 13.6778 | < 0.001 |
| | Age Class-Weaner | 1.5194 | 0.4792 | 0.5610 | 2.4778 | < 0.01 |
| | Body Length | 0.0427 | 0.0260 | -0.0093 | 0.0947 | 0.105 |
| | Body Mass | 0.1560 | 0.0587 | 0.0386 | 0.2734 | < 0.01 |
| | Sex-Male | 0.4523 | 0.3032 | -0.1541 | 1.0587 | 0.141 |

*Note.* The vocal tract structures tested are vocal tract length (VTL), vocal fold length (VFL), subglottic tracheal dorsoventral distance 1 (STDV1) and subglottic tracheal dorsoventral distance 2 (STDV2). For all models, the reference level for Age Class is ‘Pup’ and the reference level for Sex is ‘Female’.

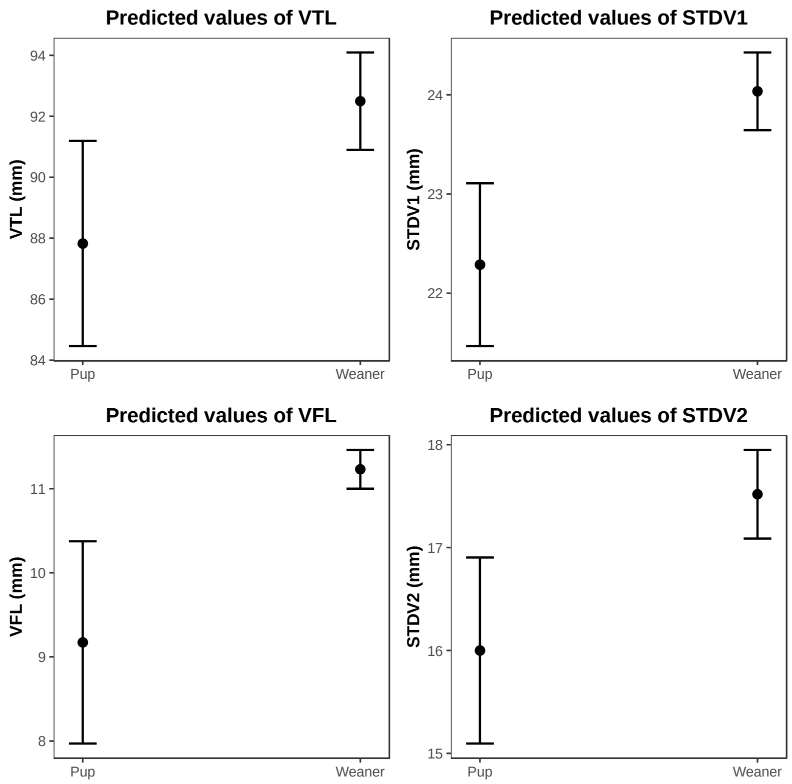


**Figure 1.** Predicted effects of Age class in each of the GLM models.

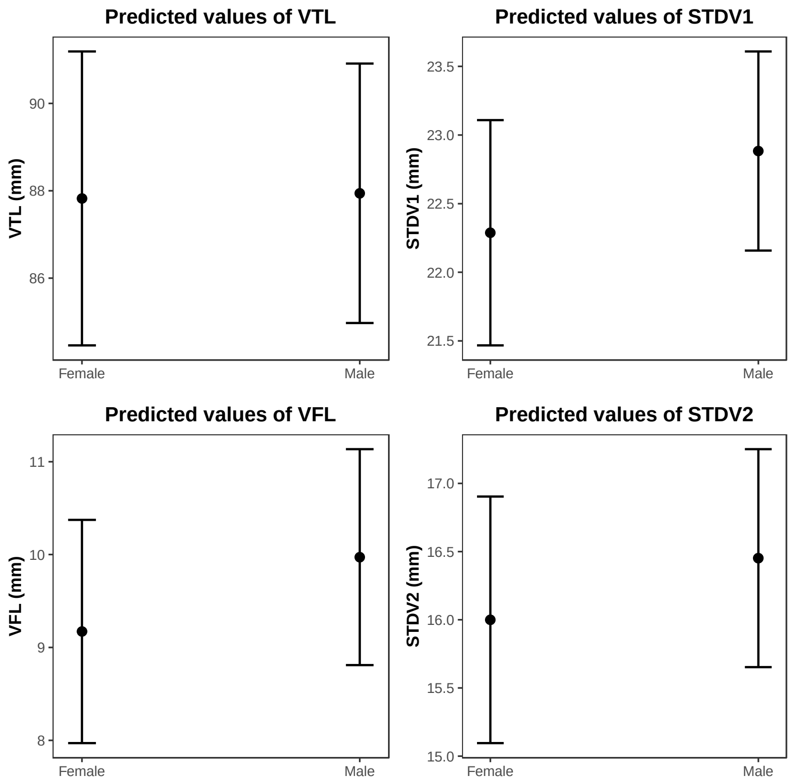


**Figure 2.** Predicted effects of Sex in each of the GLM models.

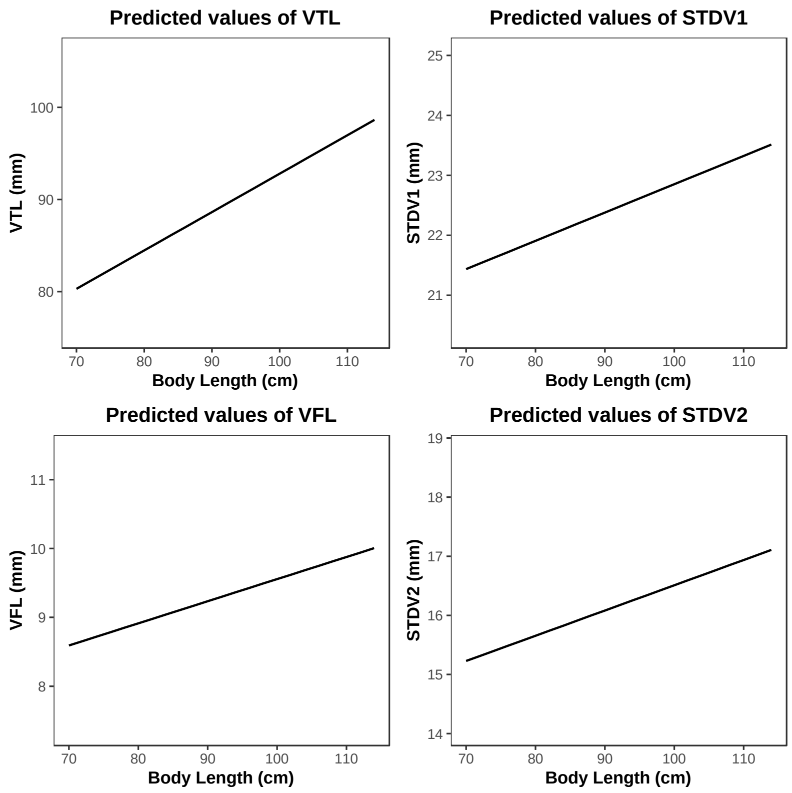


**Figure 3.** Predicted effects of Body Length in each of the GLM models.

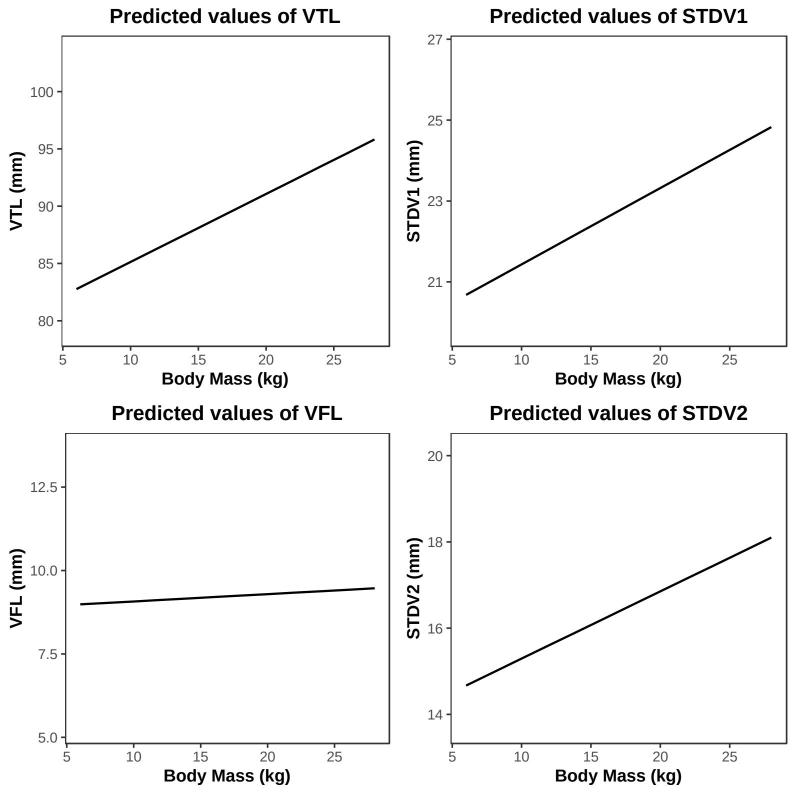


**Figure 4.** Predicted effects of Body Mass in each of the GLM models.

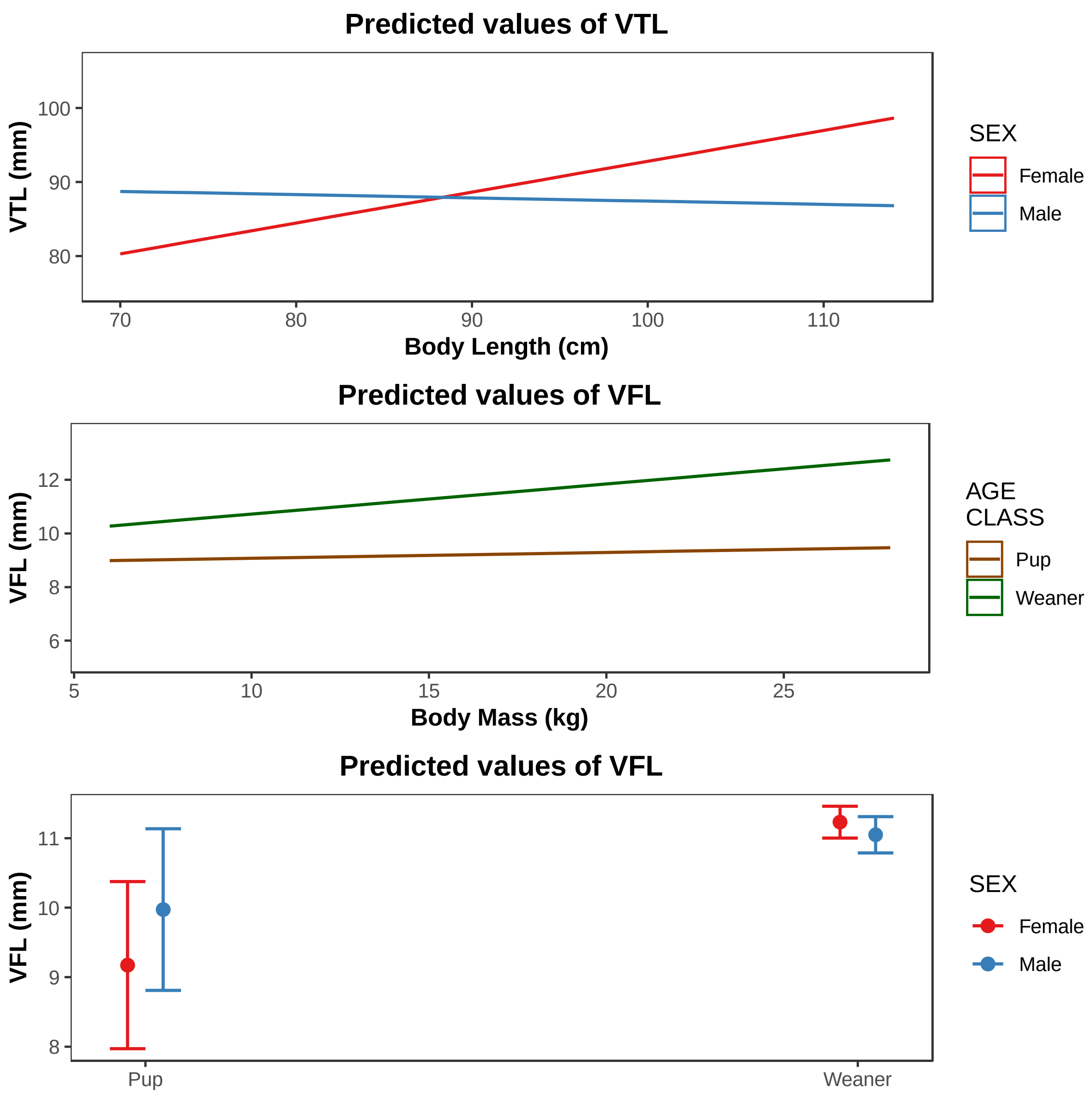


**Figure 5.** Predicted effects of the body length and sex interaction for VTL (top), the body mass and age interaction for VFL (middle) and the age and sex interaction for VFL (bottom).
